# Novel hemorrhage models of cerebral cavernous malformations

**DOI:** 10.1101/2020.02.12.944421

**Authors:** Matthew R Detter, Robert Shenkar, Christian R Benavides, Catherine A Neilson, Thomas Moore, Rhonda Lightle, Nicholas Hobson, Le Shen, Ying Cao, Romuald Girard, Dongdong Zhang, Erin Griffin, Carol J Gallione, Issam A Awad, Douglas A Marchuk

## Abstract

Cerebral cavernous malformations (CCMs) are ectatic capillary-venous malformations that develop in approximately 0.5% of the population. Patients with CCMs may develop headaches, focal neurologic deficits, seizures, and hemorrhages. While symptomatic CCMs, depending upon the anatomic location, can be surgically removed, there is currently no pharmaceutical therapy to treat CCMs. Several mouse models have been developed to better understand CCM pathogenesis and test therapeutics. The most common mouse models induce a large CCM burden that is anatomically restricted to the cerebellum and contributes to lethality in the early days of life. These inducible models thus have a relatively short period for drug administration. We developed an inducible CCM3 mouse model that develops CCMs after weaning and provides a longer period for potential therapeutic intervention. Using this new model, three recently proposed CCM therapies - fasudil, tempol, vitamin D_3_, and a combination of the three drugs failed to substantially reduce CCM formation when treatment was administered for five weeks, from postnatal day 21 (P21) to P56. We next restricted Ccm3 deletion to the brain vasculature and provided greater time (121 days) for CCMs to develop chronic hemorrhage, recapitulating the human lesions. We also developed the first model of acute CCM hemorrhage by injecting mice harboring CCMs with lipopolysaccharide. These efficient models will enable future drug studies to more precisely target clinically relevant features of CCM disease: CCM formation, chronic hemorrhage, and acute hemorrhage.

## Introduction

Cerebral cavernous malformations (CCMs), also known as cavernous angiomas, are clusters of dilated and brittle capillary-venous vessels that develop in approximately 1 in 200 individuals [1]. CCMs can develop sporadically or in an autosomal dominant pattern of inheritance. Sporadic CCM disease often presents as a solitary lesion in adulthood while the familial form typically consists of multiple CCMs with an earlier onset of disease. These distinct differences in clinical presentation of sporadic and familial CCMs led researchers to hypothesize that CCM pathogenesis follows a two-hit model of disease as originally described by Knudson [2]. The genetic basis of CCM disease was subsequently discovered to be biallelic loss-of-function mutations in one of three genes: *KRIT1 (KREV1/RAP1A interaction trapped-1)/CCM1* [3, 4], *OSM (Osmosensing scaffold for MEKK3)/CCM2* [5, 6], or *PDCD10 (Programmed cell death 10)/CCM3* [7]. Further evidence for the two-hit genetic model of CCM pathogenesis came from the identification of second somatic mutations with DNA sequencing of CCMs [8-11].

Despite the different clinical presentation of the sporadic and familial CCM diseases, sporadic and familial CCMs share the same symptomology of headaches, focal neurologic deficits, seizures, and hemorrhage. Symptomatic hemorrhage occurs in the familial disease at an approximate frequency of 6.5% per patient per year [12]. A meta-analysis of 1620 individuals with sporadic or familial CCMs observed a 15.8% five-year risk of symptomatic hemorrhage following CCM diagnosis[13]. Following a symptomatic hemorrhage, the five-year risk of a recurrent hemorrhage is 42.4% [14]. Symptomatic CCMs can be surgically resected; however, the location of the CCM and risk of associated morbidity from surgery precludes many individuals from undergoing surgical intervention. Individuals who are not candidates for surgery receive medical therapy for symptom management, but there are currently no therapies to target the etiology and bleeding sequela of CCMs.

This lack of medical therapies is not due a lack of mechanistic knowledge of CCM pathology or a lack of therapies tested in animal models. The research community has identified a number of different signaling pathways dysregulated following loss of the CCM genes: RhoA/ROCK [15-18], MEKK3-KLF2/4 [19-21], ICAP-1 and β1 integrin [22, 23], DELTA-NOTCH [24], angiopoietin-2 [25], thrombomodulin and endothelial protein C receptor [26], reactive oxygen species (ROS) [27], autophagy [28], and endothelial-to-mesenchymal transition (EndMT) [29]. A nearly equal number of therapeutics have also been proposed or tested in CCM models: statins [15], fasudil [30], TGF-β inhibitors [29], sulindac [31], tempol [32], vitamin D_3_[32], angiopoietin-2 neutralizing antibody [25], fluvastatin and zoledronate [33], indirubin-3’-monoxime [34], thrombospondin1 replacement [35], propranolol [36, 37], ponatinib [38], BA1049 [39], and VEGFR2 inhibitor [40]. Several of these compounds arose from unbiased, high-throughput *in vitro* and *in vivo* screens of libraries containing thousands of compounds [32-34]. However, despite the extensive mechanistic and therapeutic studies of CCM disease, a robust pharmacologic treatment for CCMs remains elusive.

The collective mechanistic knowledge of CCMs is derived from studies with CCM models in the worm [41], zebrafish [42-46], and mouse [47, 48, 15, 45, 49-52]. Our lab has focused on developing different mouse models of CCMs that follow the two-hit mechanism of the human disease. We developed CCM mouse models that randomly acquire a second somatic mutation in either *Krit1, Ccm2*, or *Pdcd10*. We generated these mice by breeding CCM heterozygous animals onto either *Msh2* or *Trp53* null backgrounds that lack important DNA repair mechanisms [53, 51]. The CCMs that develop in these models have the same chronic hemorrhage and inflammatory infiltrates that are central to the human disease [53, 51]. These models have been used in five-month long studies of statin and fasudil therapy [30, 54, 55]. A drawback of this model for preclinical drug testing is the stochastic nature of when the second somatic mutation occurs. The stochasticity of this model results in CCMs that closely recapitulate the human disease but develop at an unknown time and at a relatively low frequency.

An alternative to the genetically sensitized CCM mouse models is the cre/loxP inducible models [50, 52, 56-58]. These inducible models contain cell-specific cre recombinase transgenes that can be induced with tamoxifen (TMX) to delete loxP-flanked, or floxed, transgenic alleles. These models provide exquisite temporal and cell-specific control of Ccm deletion. A surprising finding replicated by many laboratories is that the induction of a phenotype in these models requires Ccm deletion in the first three days of neonatal life. With few exceptions [52, 58], the CCM phenotype of these inducible models develops within the first week of life and is restricted nearly exclusively to the cerebellum. Both the combination of CCM deletion required within the first few days of life and the anatomic restriction of the CCMs to the cerebellum has led to the hypothesis that the developmental angiogenesis continuing after birth in the cerebellum plays a significant role in the CCM phenotype of the inducible model. However, unlike the CCMs in these inducible mouse models, human CCMs are not anatomically restricted and develop during all decades of life when developmental angiogenesis has ceased. The most significant limitation of the inducible mouse models is the lethality near weaning due to a severe CCM burden, and the absence of hemorrhage and inflammation in the lesions [59]. The lethality of this model soon after CCM formation presents a challenge for administering potential therapies over an extended period of time.

An ideal CCM mouse model would combine the desirable features of both the genetically sensitized and the inducible models. This hypothetical mouse model would develop CCMs that have the following characteristics: initiate after the period of early developmental angiogenesis, develop at a known and reproducible time following Ccm deletion, exhibit a sufficient total CCM burden to measure treatment effects while not leading to early lethality, occur proportionally throughout the brain, contain inflammatory infiltrates, and develop chronic and acute hemorrhages. We have developed novel CCM3 mouse models in which CCMs containing the desirable features listed above can be induced to better model the human disease for preclinical therapeutic studies.

## Materials and Methods

### Mice

All experiments were approved by the Duke University Institutional Animal Care and Use Committee (IACUC). We generously received the following transgenic alleles from other investigators: *Pdgfb-iCreET2* [60], *Ccm3*^*Flox*^ and *Ccm3*^*KO*^ [50], and *Slco1c1(BAC)-CreERT2* [61]. These mice were maintained on a C57BL/6J background. Mice were administered a single dose of tamoxifen (Sigma T5648) in a 9:1 (vol:vol) corn oil to ethanol solution on P1 or P6 to induce CCMs. Mice were either inject with 10μg or 25μg of tamoxifen, the dose for each experiment is noted in the results section. Gross brain images were captured with a Nikon SMZ-2T dissecting scope and Leica DFC425 camera with Leica Application Suite version 3.8.0 software. The yellow contrast of the entire image in Figure 1 B was auto adjusted in ImageJ to appear consistent with the other gross brain images. No other images were adjusted.

**Figure 1.**
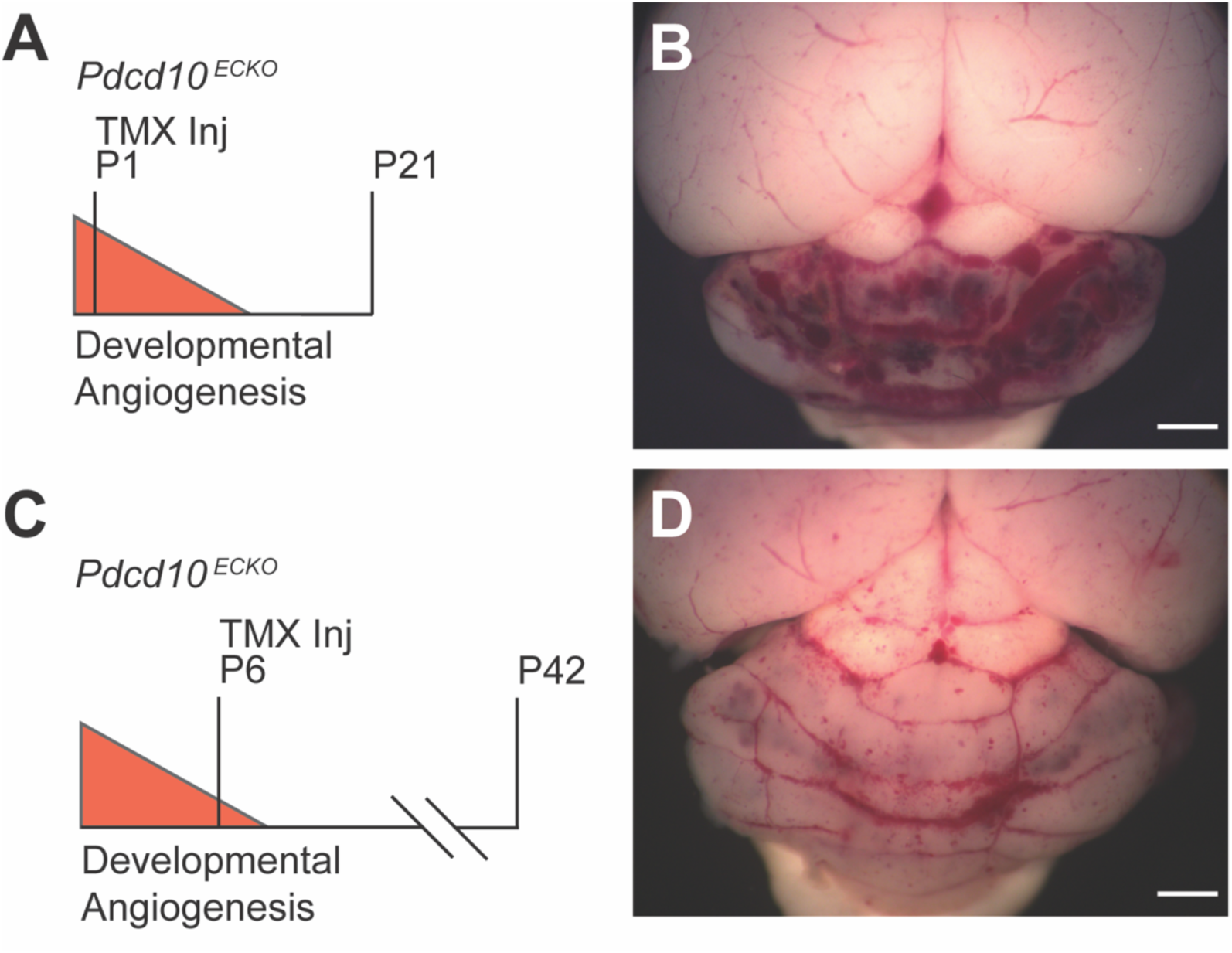
Delaying *Pdcd10* deletion resulted in a reduced CCM burden and improved viability. A) The traditional strategy of deleting *Pdcd10* on postnatal day 1 (P1) results in B) a large CCM burden restricted to the cerebellum and lethality near P21 as shown in a representative mouse. C) Delayed *Pdcd10* deletion on P6 results in D) a moderate CCM phenotype and improved viability to at least P42 as shown in a representative mouse. (scale bars: 1 mm)

### Fasudil, tempol, and vitamin D_3_ studies

We used fasudil (100 mg/kg/day in the drinking water, LC Laboratories F-4660), tempol (170 mg/kg/day in the drinking water, Sigma-Aldrich 176141), vitamin D_3_ (25 IU/g in the chow, Envigo Teklad diet, Vit D3 Suppl Diet, TD.110800) and a combination of these three drugs at the same doses for the monotherapy and triple therapy drug studies. Prior to the studies, we monitored the consumption of water by mice given fasudil, tempol, and a combination of fasudil and tempol at the experimental doses. We did not observe any differences in the amount of water consumed or in body weights of these pilot groups when compared to vehicle treated mice. Experimenters were blinded to the genotype of the pups at the time of tamoxifen injection as well as blinded to the CCM burden of the mice when enrolled in treatment groups. In an effort to reduce the effect of litter-to-litter variability, we enrolled mice from each litter into the vehicle and at least one of the treatment groups. This enrollment pattern ensured that the comparison of a treatment group and the vehicle was primarily between littermates, as opposed to entire litters enrolled in a single group and the comparison being primarily across litters. This enrollment pattern also resulted in a greater number of mice in the vehicle group than any of the monotherapy groups. Prior to the triple therapy study, we planned to compare the CCM burden of the vehicle group in the monotherapy study to the vehicle group in the triple therapy study and combine the two groups for a more balanced statistical analysis in the triple therapy study if there was no statistical differences between the two vehicle groups. Therefore, we randomly assigned more mice into the treatment group than the vehicle group of the triple therapy study. A total of 176 mice were enrolled into the monotherapy and triple therapy studies with approximately equal numbers of male and female mice in each group. The mice were inspected daily for well-being and weekly body weights were collected to track growth and overall health. There were 11 animals that underwent attrition from the following groups: 5 vehicle, 2 fasudil, 1 tempol, 2 vitamin D_3_, and 1 triple therapy.

Researchers at the University of Chicago who performed the microCT analysis were blinded to the treatment group assignments during data gathering and were unblinded only after all brain images had been processed. Lesion volume normalized to total brain volume was determined by micro-computed tomography (microCT) as previously described [39, 55, 59, 51]. A 1-mm thick coronal slice that included the most abnormalities, as observed with micro-CT, was cut with a mouse brain matrix. The slice was processed, embedded in paraffin, and cut into 5-μm thick sections with a microtome. The sections were placed onto microscope slides and stained as previously reported with hematoxylin and eosin, Perls’ Prussian blue for non-heme iron, and anti-CD45R/B220 antibody for B lymphocytes. To visualize endothelium, sections were incubated with 5 μg/ml goat anti-mouse CD31 (R&D Systems, Inc, Minneapolis, MN) overnight at 4°C, followed by 1:1000 donkey anti goat secondary antibody conjugated with Alexa Fluor 647 (Jackson ImmunoResearch Laboratories, Inc, West Grove, PA) for 1 h at room temperature, and observed under Axiover 200M (Carl Zeiss AG, Oberkochen, Germany) microscopy. Red blood cell auto-fluorescence was detected at approximately 580 nm. Image files were exported as grey scale TIF images, and converted to RGB format by using Image J (National Institutes of Health, Bethesda, MD) as described previously.[59]

### Lipopolysaccharide (LPS) experiments

LPS from Escherichia coli O111:B4 (Sigma L2630) was dissolved in sterile PBS at a concentration of 0.1mg/mL. LPS was administered at a dose of 250 μg/kg via intraperitoneal injection. Thus, a 20g mouse would receive a 50-μL injection of 0.1mg/mL LPS stock solution for a dose of 250μg/kg. Mice that lost >20% of their starting body weight were euthanized per the humane endpoints of our study.

### Statistical Analysis

Statistical outliers in the lesion burden data from the microCT quantification were defined as >2 standard deviations above or below the mean of their respective group (Z-score > 2). Six mice were removed from the analysis as outliers from the following groups: 1 vehicle, 1 fasudil, 2 tempol, 1 vitamin D_3_, and 1 triple therapy. The final number of mice in each group were: 31 vehicle from the monotherapy study, 20 fasudil, 22 tempol, 22 vitamin D_3_, 23 additional vehicle from the triple therapy study, and 40 triple therapy. All graphs are dot plots with the individual values and mean ± standard error of the mean (SEM) shown. The assumptions of normality and homogeneity of variances were tested with D’Agostino & Pearson omnibus normality and Levene’s tests, respectively. Statistical significance (p < 0.05) was calculated with one-way ANOVA and post hoc Bonferroni tests, independent samples two-tailed t test, and independent samples Mann-Whitney U test. The specific test for each analysis is listed in the figure legends. Statistical analysis was performed with SPSS version 26. Graphs were generated with GraphPad Prism 8.

## Results

### Delaying Pdcd10 deletion delays CCM formation and extends the viability of the inducible CCM3 mouse model

We investigated two strategies to extend the viability of the inducible CCM3 mouse model: 1) decreasing the dose of tamoxifen (TMX) administered to activate the cre recombinase and 2) inducing *Pdcd10* deletion later in the postnatal period. Since their development, the inducible CCM alleles have required deletion during the first three days of life to induce CCMs. We hypothesized that deleting *Pdcd10* later in the period of developmental angiogenesis may induce a reduced CCM burden that is conducive to drug studies lasting several weeks. We administered 10μg of TMX to *Pdcd10*^*Flox/Flox*^ [50], *Pdgfb-iCreERT2* [60] (hereafter abbreviated as *Pdcd10*^*ECKO*^) mice on P1 and observed the same phenotype that has been reported previously – the vast majority of CCMs in the cerebellum with lethality before or near P21 (Figure 1 A, B). Further reducing the dose of TMX on P1 did not substantially change the CCM phenotype, especially with regard to the anatomic restriction of the lesions. A more significant change in phenotype was observed when we delayed *Pdcd10* deletion to P6 in the *Pdcd10*^*ECKO*^ mice. Delaying *Pdcd10* deletion reduced the CCM burden in the cerebellum and extended the viability to at least P42 (Figure 1 C, D). This reduced CCM burden following deletion of *Pdcd10* later in the developmental angiogenic window suggested that we could create a model with a sufficient CCM burden and viability to perform therapeutic studies over several weeks.

We identified a novel *Pdcd10* deletion strategy that resulted in a progressive CCM phenotype beginning after weaning. We injected *Pdcd10*^*ECKO*^ mice with 25μg of TMX on P6. The first CCMs in these *Pdcd10*^*ECKO*^ mice were visible to the eye at approximately P28 (Figure 2 A), well after the developmental angiogenesis. To determine the full extent of the lesion burden and the anatomic distribution of lesions across the entire brain, we used micro computed tomography (microCT). Precise CCM volumes were measured and reported as a percentage of total brain volume (Figure 2 B). Gross images of the dorsal and ventral surfaces of the brain along with 3D renderings of the microCT scans demonstrated a progressive CCM burden that began in the cerebellum and later involved the entire brain (Figure 2 C-K). Histology determined that the CCMs in this model are predominately single cavern (stage 1) lesions without an appreciable amount of multicavernous (stage 2) lesions. Observing primarily vascular ectasia lesions (dilated individual caverns) at all three timepoints (P35, P42, and P70), rather than complex multicavernous lesions, suggested that the increase in CCM burden over time was due to the formation of new CCMs rather than an increase in size of the first CCMs to develop. The strength of the *Pdcd10*^*ECKO*^ model was the continued formation of CCMs over a 40-day period.

**Figure 2.**
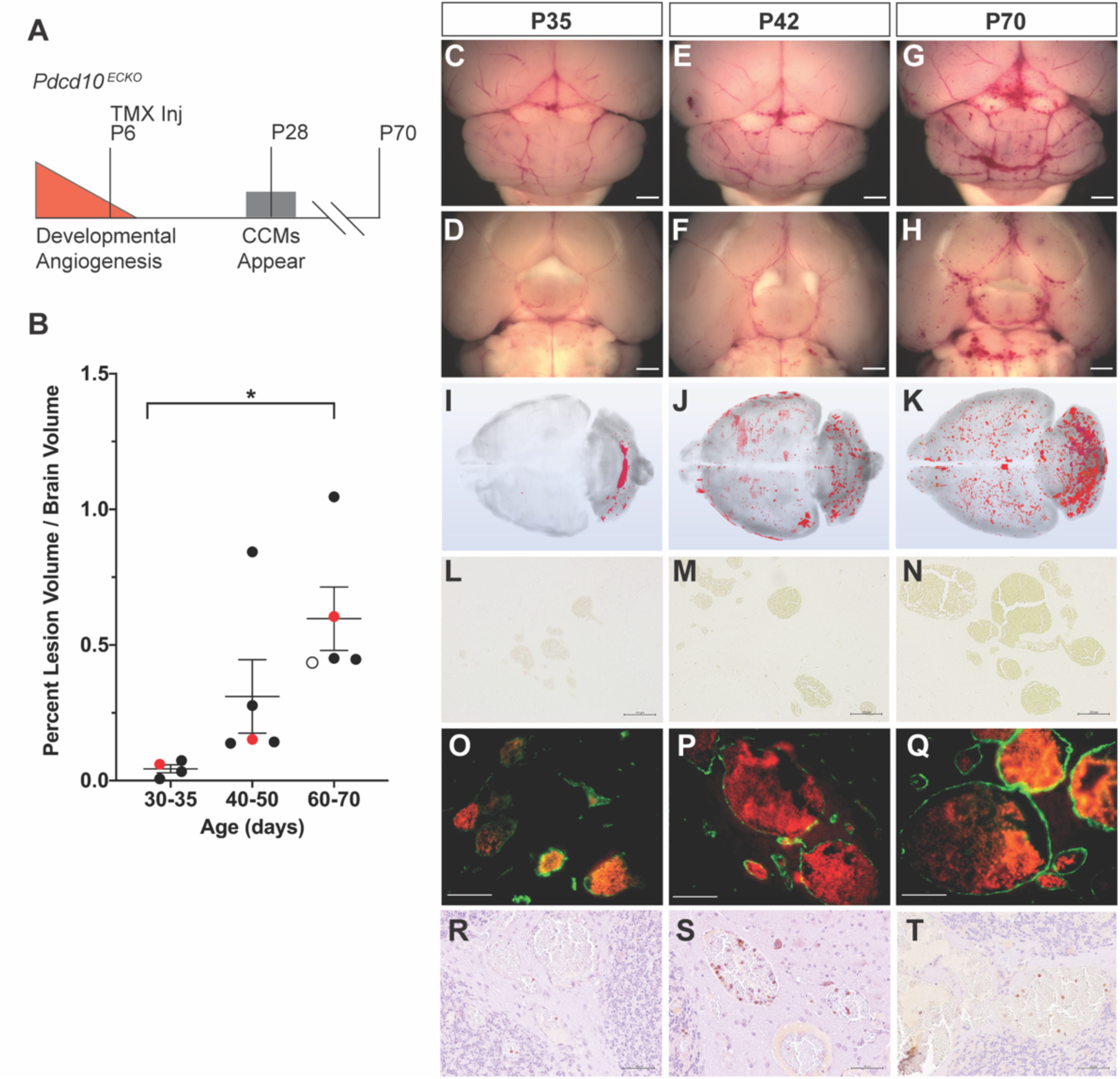
*Pdcd10* deletion on P6 results in a progressive phenotype that develops new CCMs through P70. A) Deletion of *Pdcd10* on P6, late in the developmental angiogenic period, results in the first CCMs appearing near P28 and lethality at approximately P70. B) MicroCT quantification of the progressive CCM burden. The red data points on the graph correspond to the P35, P42, and P70 mice characterized in the subsequent panels in the figure. The open circle data point in the P60-70 groups corresponds to an attrition mouse shown in Supplemental Figure 1. Data are presented as each value, the mean, and standard error of the mean. Statistical significance was determined with one-way ANOVA (p=0.018) and post hoc Bonferroni test, p=0.017. C-H) Gross images of the dorsal and ventral surfaces (scale bars: 1mm) and I-K) 3D renderings of the microCT scans demonstrate the progression of CCMs from the cerebellum at P35 to the entire brain by P70. L-N) Perls’ Prussian blue staining demonstrates primarily single cavern, stage 1, lesions with a paucity of chronic hemorrhage (scale bars: 100μm). O-Q) Endothelium stained with anti-CD31 (green) and red blood cell autofluorescence within the lumen of the CCMs demonstrates the lack of acute hemorrhage into the brain parenchyma in this model (scale bars: 50μm). R-T) B lymphocytic infiltrates were observed in CCMs at P42 and P70 (scale bars: 50μm).

The improved viability of the *Pdcd10*^*ECKO*^ model enabled the CCMs to acquire immune cell infiltrates over time. Lymphocytic infiltrates are an important component of the human disease [62] and depletion of B lymphocytes in a genetically sensitized CCM3 mouse model reduced the formation of stage 2 lesions and chronic hemorrhage [63]. A subpopulation of CCMs within the *Pdcd10*^*ECKO*^ model contained B cell infiltrates beginning at approximately P42 (Figure 2 R-T). The presence of B lymphocytes in the *Pdcd10*^*ECKO*^ model indicated that this inducible model contained CCMs that recapitulated important pathologic features of the human disease. As expected for a model exhibiting primarily single caverns lesions, CCMs in the *Pdcd10*^*ECKO*^ model did not exhibit significant amounts of chronic or acute hemorrhage. Chronic hemorrhage was visualized with Perls’ Prussian blue dye that stained non-heme iron surrounding the malformations (Figure 2 L-N). Acute hemorrhage was measured by staining the CCM endothelium with CD31 and imaging the autofluorescence of red blood cells extravasated from the malformation lumen and into the surrounding brain parenchyma (Figure 2 O-Q). The induced CCMs within the *Pdcd10*^*ECKO*^ mouse began to exhibit characteristics of the human pathology, namely the presence of B lymphocytes, but lacked the hemorrhage phenotype.

The limit of viability for the *Pdcd10*^*ECKO*^ model was approximately 70 days. Subsequent to P70, we observed systemic pathologies, particularly within the gastrointestinal tract. Gross examination of the internal organs of a mouse that required early euthanasia revealed ischemic bowels (Figure 2B open circle, Supplemental Figure 1). Histology of the small and large intestines identified several abnormalities in the distal colon: dilated lamina propria micro-vessels, crypt abscesses, and epithelial erosion with granulation tissue formation (Supplementary Figure 1). The colon abnormalities are consistent with previous studies of Krit1 [64-66] and Pdcd10 [67] in intestinal epithelial barrier function and gastrointestinal pathologies. The fundamental roles of CCM proteins in intestinal cells is conserved across species as they have been observed in *C. elegans* [66] and mouse [67] models of CCM. Despite the development of severe gastrointestinal disease by P70, the extended life span over acute, perinatal models of CCM disease enabled the use of the *Pdcd10*^*ECKO*^ model to test the ability of small molecule therapeutics to block the formation of CCMs.

### Neither fasudil, tempol, vitamin D_3_, nor triple therapy substantially reduced CCM formation with five weeks of treatment

We tested three of the more promising proposed therapies for CCM disease, fasudil, tempol, and vitamin D_3_, in the novel *Pdcd10*^*ECKO*^ model. Fasudil, tempol, and vitamin D_3_ have each demonstrated modest effect in reducing CCM formation in either CCM1 [30, 54], CCM2 [32] or CCM3 [55] mouse models. We previously reduced the CCM burden of genetically sensitized CCM1 and CCM3 mouse models through the inhibition of the Rho-kinase signaling pathway with fasudil [30, 54, 55]. We selected fasudil for the current study because thus far, it had passed the test of reproducibility (different studies) and generality (different genetic models) for CCM disease. Tempol and vitamin D_3_ were chosen because they were the top two candidates from an *in vitro* screen of over 2,000 compounds; and subsequently, both tempol and vitamin D_3_ demonstrated efficacy in reducing the CCM burden in an inducible CCM2 mouse model [32]. Tempol is a superoxide dismutase mimetic with intracellular antioxidant activity that reduces reactive oxygen species. The primary signaling pathways by which vitamin D_3_ improved CCM disease is not currently known [32]. Thus far, neither tempol nor vitamin D_3_ had been tested in a CCM3 model. Because fasudil, tempol, and vitamin D_3_ have shown only moderate effects when administered as monotherapies, we also combined all three drugs to discover any potential additive or synergistic effects of combination therapy. Although combination therapy to more fully inhibit a single pathway has been attempted [33], combination therapy targeting multiple, distinct molecular pathways has never been tested in CCM disease.

We randomly assigned mice to vehicle (n=36), fasudil (n=23), tempol (n=25), and vitamin D_3_ (n=25) groups for five weeks of treatment from P21 to P56 (Figure 3 A). We calculated a sample size of 22 mice for each group to achieve 90% power (alpha 0.05) to see an effect. We chose our minimal effect to be the same difference in lesion burden between mice at P60-70 and P40-50 (a 48% difference) (Figure 2 B). Fasudil (100 mg/kg/day in the drinking water) and vitamin D_3_ (25 IU/g in the chow) we administered at the same dose as previous CCM studies [30, 32]. We doubled the dose of tempol (172 mg/kg/day in the drinking water) reported in a previous CCM study [32] based on reports of higher doses being well tolerated and our pre-study monitoring of drug consumption (see Materials and Methods). We concluded the study at P56 so as to minimize the number of attrition animals lost to systemic pathologies as described above. Nonetheless, concluding the study at P56 provided a 5-week period after weaning at P21 for a sufficient CCM burden to develop. The total CCM volume (mm^3^) throughout the entire brain was measured with microCT and reported as a percentage of total brain volume. Researchers performing the microCT analysis were blinded to the treatment group assignments. Fasudil (p=0.058) and tempol (p=0.048) exhibited decreased CCM lesion burden; however, we did not observe a substantial decrease in CCM burden following five weeks of treatment with fasudil, tempol, or vitamin D_3_ monotherapies in the *Pdcd10*^*ECKO*^ mice (Figure 3 B).

**Figure 3.**
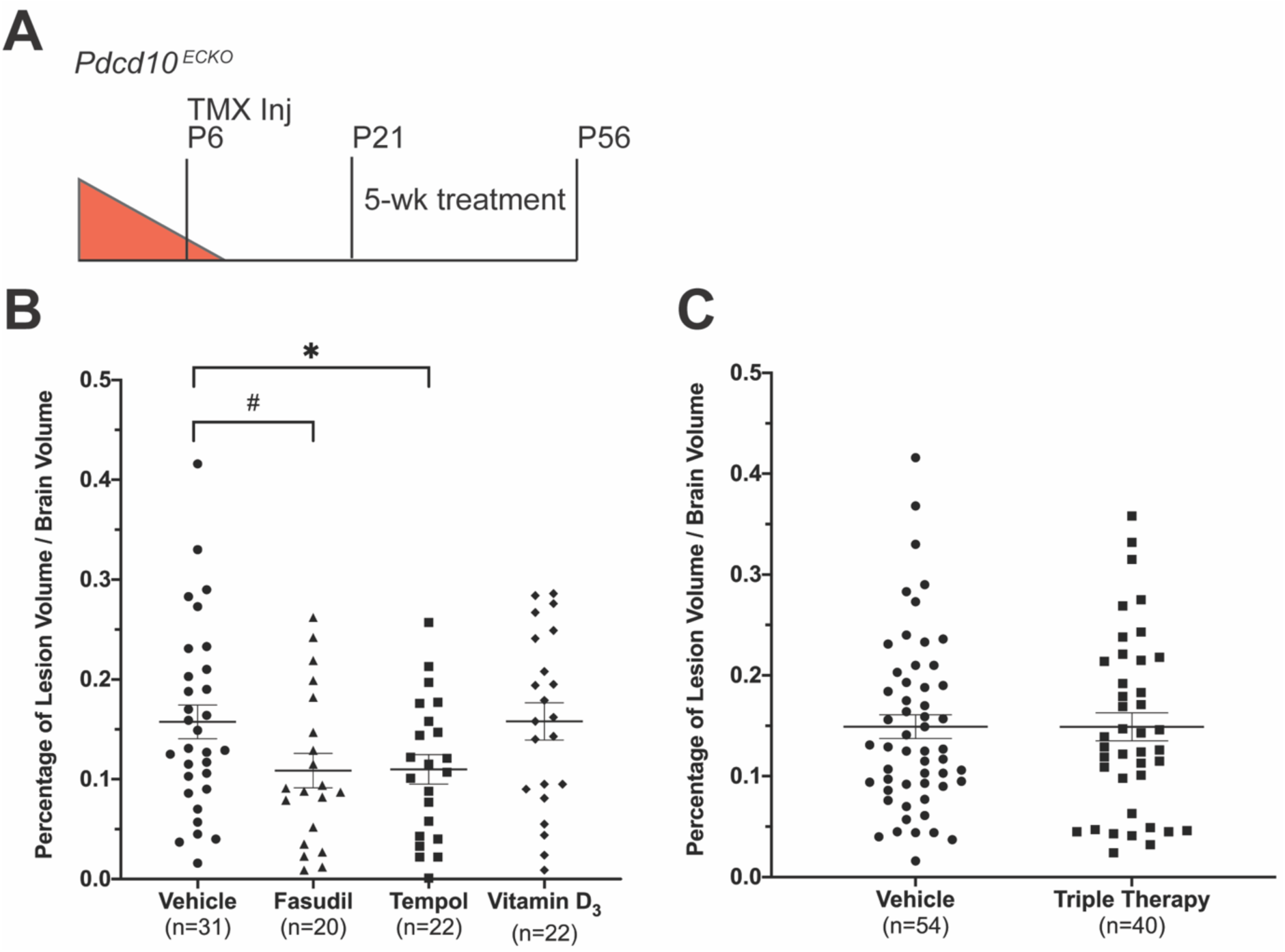
Five weeks of fasudil, tempol, vitamin D_3_, or triple therapy treatment does not substantially reduce CCM formation in the *Pdcd10*^*ECKO*^ model. A) We designed the studies with *Pdcd10* deletion on P6 followed by 5 weeks of drug treatment from P21 to P56. Total CCM volume was measured by microCT and reported as percentage of total brain volume. Data are presented as each value, the mean, and standard error of the mean. B) Lesion burden in the monotherapy study was reduced with fasudil (a trend with p=0.058 (#), n=20) and tempol (statistically significant p=0.048 (*), n=22), but not with vitamin D_3_ (p =0.987, n=22) when compared to vehicle (n=31). Statistical significance was determined by the independent samples two-tailed t test. C) The triple therapy study with vehicle (n=54) and fasudil, tempol, and vitamin D_3_ combination treatment (n=40) did not reach the level of statistical significance as determined by the Mann-Whitney U test, p=0.878.

The trend of decreased CCM burden following fasudil and tempol monotherapy suggested that additive or synergistic therapeutic effects may occur with triple therapy of fasudil, tempol, and vitamin D_3_. We conducted a second study comparing vehicle (n=24 additional mice) and triple therapy (n=41). The triple therapy combined each of the monotherapy drugs at the same respective doses as the previous study. To perform a more balanced statistical analysis of the triple therapy study, we added the vehicle treated mice from the monotherapy study with the vehicle treated mice in the triple therapy study (n=54) (Figure 3 C). The CCM burden of the vehicle treated mice from the monotherapy and triple therapy studies were not statistically different. The mice treated with triple therapy did not show any reduction in CCM formation (Figure 3 C). The modest effects seen in the monotherapy study did not translate to additive or synergistic effects in the triple therapy study. The results of the triple therapy study reinforce the findings of the monotherapy study that neither fasudil, tempol, nor vitamin D_3_ can substantially reduce CCM formation in the *Pdcd10*^*ECKO*^ model. This was the first study to test a combination of drugs that target distinct molecular pathways of CCM pathology. The inability of fasudil, tempol, and vitamin D_3_ to reduce CCM formation in the model highlights both the potential differences in CCM response to therapies depending upon the underlying genetic mutation and the need for continued research to identify a more robust medical therapy that impacts lesion development.

### Restricting Pdcd10 deletion to the cerebral vasculature enables CCMs to develop chronic hemorrhage

CCM formation is an early stage of the human disease. Later, clinically relevant stages of disease are chronic and acute hemorrhage. Chronic and acute hemorrhage were not observed at sufficient frequencies in the *Pdcd10*^*ECKO*^ model to serve as quantitative endpoints in the drug studies. We attributed the lack of CCM hemorrhage in *Pdcd10*^*ECKO*^ model to the relatively short period, of approximately 40 days, between the formation of the first CCMs at P28 and lethality due to systemic pathologies near P70 (Figure 2 A). We hypothesized that we could induce hemorrhagic CCMs by allowing the CCMs to mature over a longer period of time. To eliminate the systemic disease that hindered the long-term viability of the *Pdcd10*^*ECKO*^ model, we replaced the global endothelial cell-specific cre recombinase (*Pdgfb-iCreERT2*) [60] with a brain endothelial cell-specific cre recombinase (*Slco1c1(BAC)-CreERT2*) [61]. The *Slco1c1(BAC)-CreERT2* transgene restricted *Pdcd10* deletion to the cerebral vasculature and eliminated the systemic endothelial deletion of *Pdcd10* and resulting vascular abnormalities in other organs.

We injected *Pdcd10*^*Flox/Flox*^, *Slco1c1(BAC)-CreERT2* (hereafter abbreviated as *Pdcd10*^*BECKO*^) mice with 25μg of TMX on P6 and aged the mice to 4 months (P121) (Figure 4 A). The CCM burden of the *Pdcd10*^*BECKO*^ model at P121 was more than 6.5-fold greater than the *Pdcd10*^*ECKO*^ model at P56, the end point of the drug studies (Figure 4 B). We observed a different CCM distribution pattern in the *Pdcd10*^*BECKO*^ model when compared to the *Pdcd10*^*ECKO*^ model (Figure 4 C-E). The CCMs induced in the *Pdcd10*^*BECKO*^ model did not appear to show an anatomic preference for the cerebellum but were more proportionally distributed throughout the brain. This CCM pattern more faithfully recapitulated the human disease. The most significant difference between the *Pdcd10*^*BECKO*^ and *Pdcd10*^*ECKO*^ models was the presence of chronic hemorrhage in the *Pdcd10*^*BECKO*^ model (Figure 4 F). Chronic hemorrhage was visualized with Perls’ Prussian blue dye that stained the non-heme iron surrounding the CCMs (Figure 4 F). The *Pdcd10*^*BECKO*^ model contained a mix of hemorrhagic and non-hemorrhagic CCMs. This mixture of CCMs with different hemorrhagic states also models the human disease presentation. Like the *Pdcd10*^*ECKO*^ model, the *Pdcd10*^*BECKO*^ model also contained lymphocytic infiltrates, another characteristic of human CCMs (Figure 4 G). This novel *Pdcd10*^*BECKO*^ mouse model restricts *Pdcd10* deletion to the cerebral vasculature to induce CCMs that more accurately model the human disease. The induced CCMs are more evenly distributed throughout the brain and a subset of the CCMs exhibit chronic hemorrhage and inflammation. This inducible CCM3 model is uniquely positioned to test the ability of a therapy to reduce chronic hemorrhage, a cardinal feature of the human disease.

**Figure 4.**
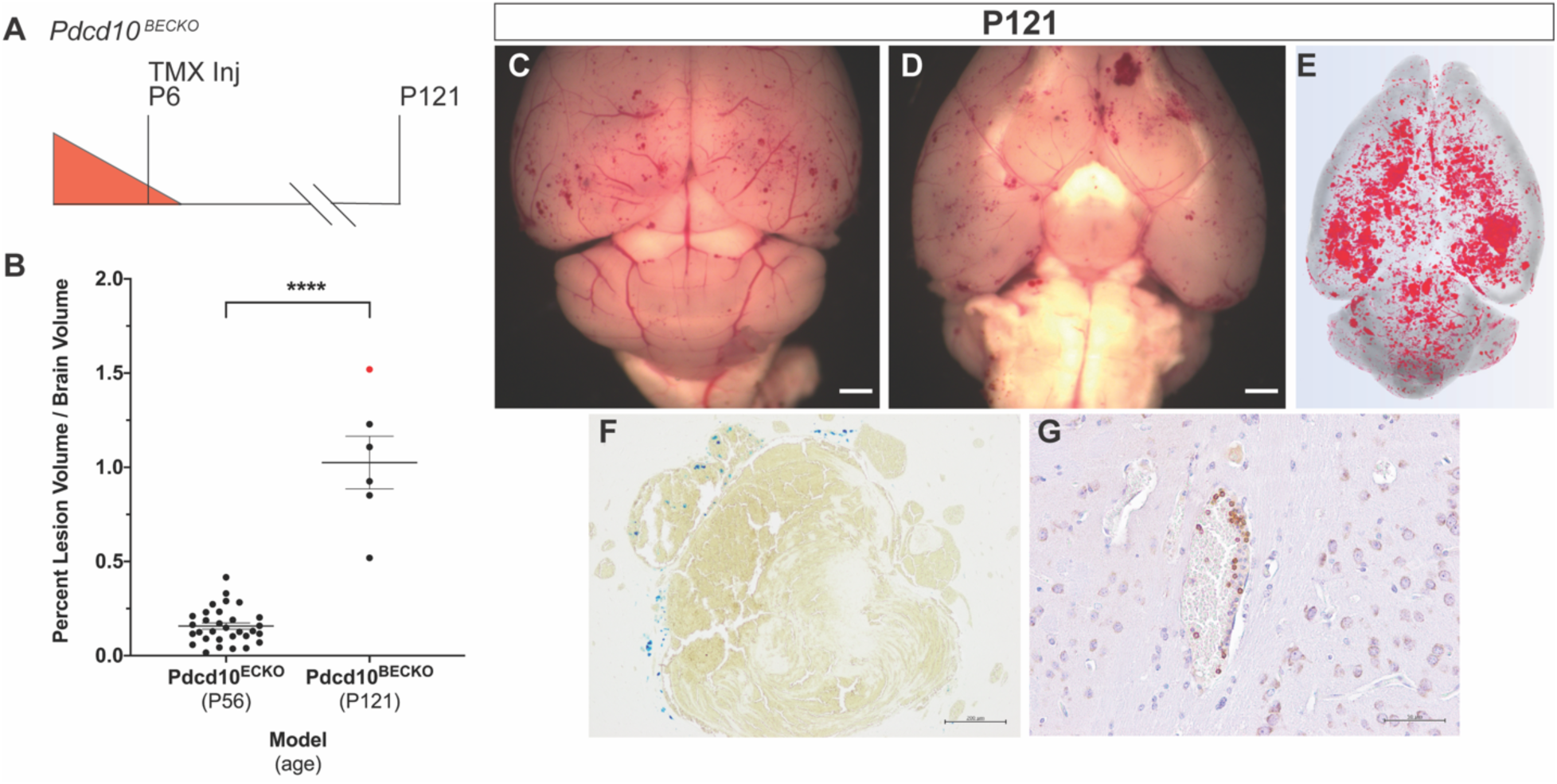
Restricting *Pdcd10* deletion to the brain vasculature (*Pdcd10*^*BECKO*^) enables development of multicavernous CCMs with chronic hemorrhage. A) Restricting *Pdcd10* deletion on P6 to the brain vasculature improves viability to at least P121 and B) leads to a 6.5-fold increase in CCM burden when compared to the *Pdcd10*^*ECKO*^ model at P56 (endpoint of the drug studies). The red data point corresponds to the individual mouse characterized in the subsequent panels in the figure. Statistical significance was determined by an independent samples Mann-Whitney U test, p<0.0001. C,D) Images of the dorsal and ventral brain surfaces along with E) microCT 3D rendering demonstrate a proportional distribution of CCMs throughout the brain (scale bars: 1mm). A subset of CCMs showed evidence of F) chronic hemorrhage, visualized with Perls’ Prussian blue stain, and G) B lymphocytic infiltrates (scale bars: F: 200μm, G: 50μm).

### Lipopolysaccharide induces acute hemorrhage in existing CCMs

Although the *Pdcd10*^*BECKO*^ mouse model exhibits chronic hemorrhage, a very important clinical feature of the human disease is acute hemorrhage. Acute CCM hemorrhages account for significant morbidity and mortality of patients. We hypothesized that acute CCM hemorrhage could be induced with an environmental stimulus. Tang et al. recently discovered a connection between the gut microbiome and CCMs that is driven by lipopolysaccharide (LPS) [21]. LPS is an endotoxin found in the outer membrane of Gram-negative bacteria. LPS signals through toll-like receptor 4 on brain endothelial cells to activate the MEKK3-KLF2/4 signaling cascade [21]. MEKK3-KLF2/4 signaling is negatively regulated by a CCM protein complex and has been demonstrated by multiple groups to be a critical pathway in CCM pathology [19, 20]. Building upon the microbiome-CCM discovery, we hypothesized that a single, sub-lethal dose of LPS may exacerbate existing CCMs and induce acute hemorrhage.

Our attempt to model acute hemorrhage in CCM was generated on a genetic background where one allele of *Pdcd10* was already deleted, thus modeling the genotype of the inherited form of CCM disease caused by autosomal dominant germline mutations in *Pdcd10*. We again restricted *Pdcd10* deletion to the brain vasculature and injected *Pdcd10*^*Flox/KO*^, *Slco1c1(BAC)-CreERT2* (hereafter called *Pdcd10*^*BECKO/KO*^) mice with 25μg of TMX on P6 to induce CCMs (Figure 5 A). We performed several pilot studies to identify an appropriate dose of LPS that resulted in an acute inflammatory response from which the mice could recover (Supplemental Figure 2). We administered LPS (250 μg/kg) via intra-peritoneal injection on P41 and collected the brains 24 hours later to analyze primarily for acute hemorrhage (Figure 5 A).

**Figure 5.**
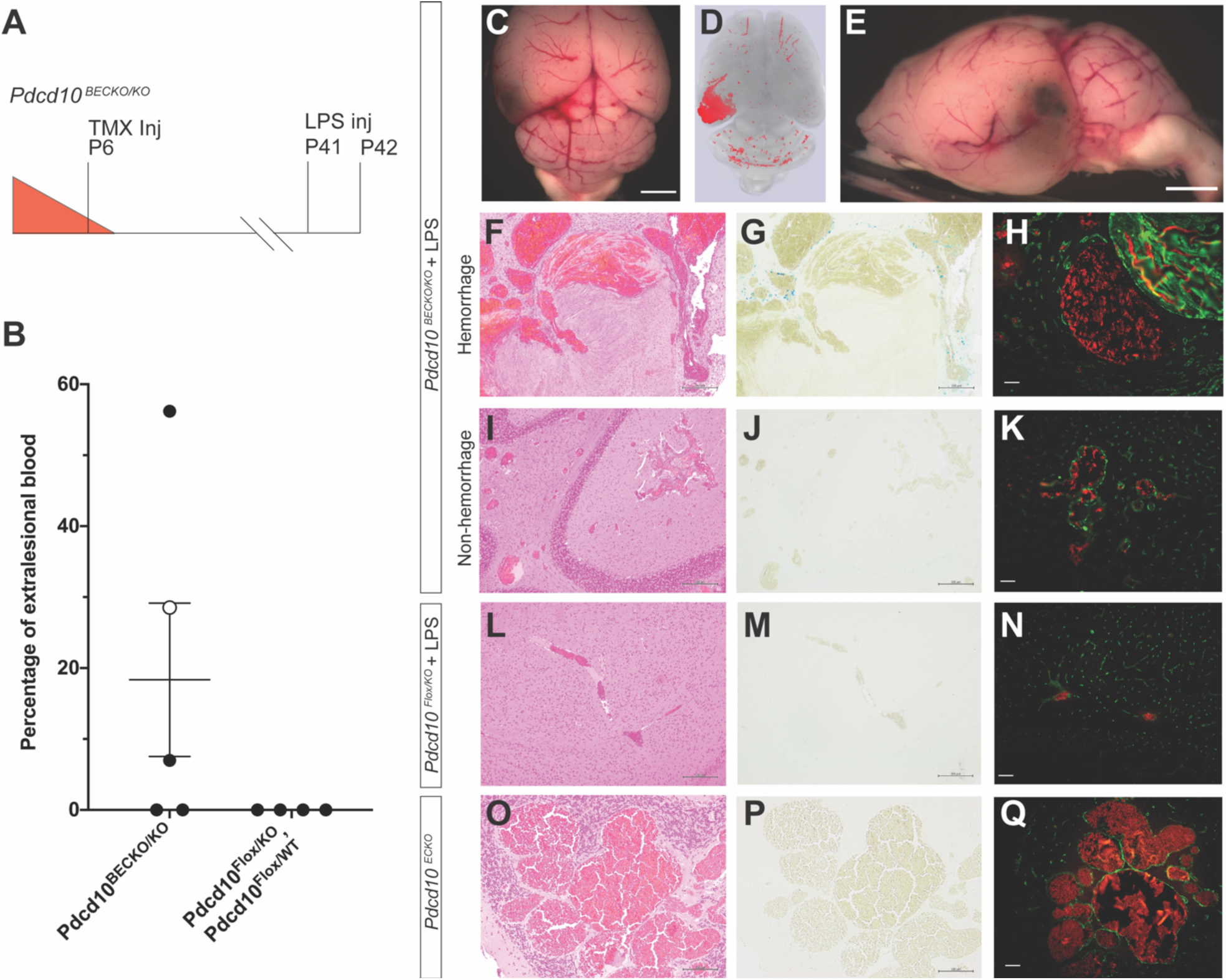
A single, sub-lethal dose of lipopolysaccharide (LPS) induces acute hemorrhage in existing CCMs. A) Restricting *Pdcd10* deletion on P6 to the brain vasculature followed by LPS injection on P41 leads to acute CCM hemorrhage within 24 hours. B) Acute hemorrhage detected in 3 of 5 LPS injected mice quantified as percentage of extralesional blood per total lesion volume. The open circle data point in the *Pdcd10*^*BECKO/KO*^ group corresponds to a mouse that was found dead and whose full characterization is shown in Supplemental Figure 3. The percentage of extralesional blood between the two groups did not reach the level of statistical significance as determined by an independent samples two-sided t test (p=0.165). C-E) Gross and microCT images demonstrating focal area of hemorrhage in the left cerebral hemisphere (scale bars: 1mm). A hemorrhagic brain region of a *Pdcd10*^*BECKO/KO*^ mouse injected with LPS was stained with F) hematoxylin and eosin (H&E), G) Perls’ Prussian blue dye, and H) CD31 labeling (green) of endothelium to visualize CCM morphology, chronic hemorrhage, and acute hemorrhage, respectively. H) Acute hemorrhage was measured from the autofluorescence of red blood cells in the brain parenchyma surrounding the CCMs. The same histologic analysis, of I) H&E, J) Perls’ Prussian blue dye, and K) CD31 labeling, was performed with samples from a nonhemorrhagic brain region of a *Pdcd10*^*BECKO/KO*^ mouse injected with LPS. K) No evidence of acute hemorrhage was detected as red blood cell autofluorescence remained within the CCM lumen. *Pdcd10*^*Flox/KO*^ mouse injected with LPS did not show formation of CCMs or extravasation of red blood cells as measured by L) H&E, M) Perls’ Prussian Blue dye, and N) CD31 labeling of the endothelium. *Pdcd10*^*ECKO*^ mice at P56 did not show evidence of extralesional red blood cells as measured by O) H&E, P) Perls’ Prussian blue dye, and Q) CD31 labeling of the endothelium. Scale bars for H&E: F-L) 200μm; O) 100μm, Perls’ Prussian Blue dye: G-M) 200μm; P) 100μm, and CD31 labeling: 50μm.

**Figure 6.**
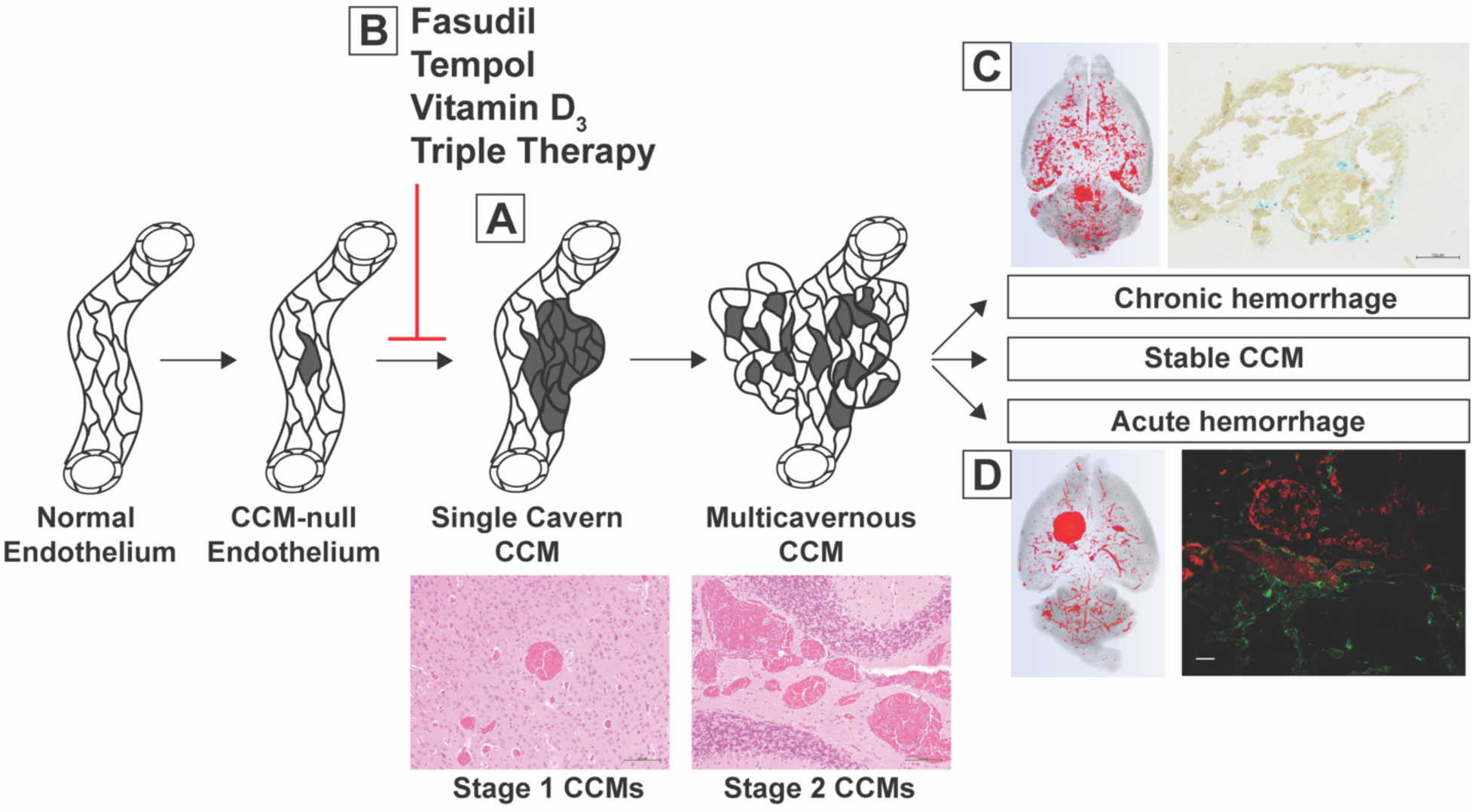
Novel CCM3 models for distinct stages of CCM disease: delayed CCM formation, chronic CCM hemorrhage, and acute CCM hemorrhage. A) We delayed *Pdcd10* deletion until P6 to induce a novel phenotype of delayed formation of single cavern CCMs (scale bars: 100 μm). B) We treated mice with this delayed phenotype for 5 weeks with fasudil, tempol, vitamin D_3_, and triple therapy combining each of the aforementioned drugs and did not detect a substantial reduction in CCM formation. C) We induced CCMs with chronic hemorrhage by restricting *Pdcd10* deletion to the brain vasculature and aging the mice to P121. A subset of multicavernous, stage 2, CCMs exhibited evidence of chronic hemorrhage as visualized with Perls’ Prussian blue dye (scale bar: 200μm). D) We induced acute CCM hemorrhage by injecting mice with existing CCMs with a single, sub-lethal dose of lipopolysaccharide. Acute hemorrhage was visualized by the autofluorescence of extravasated red blood cells in the brain parenchyma surrounding CD31 labeled CCM endothelium (scale bar: 50μm). These inducible mouse models are novel tools that will enable researchers to target specific aspects of CCM pathology in preclinical therapeutic studies.

MicroCT quantification of the LPS injected *Pdcd10*^*BECKO/KO*^ mice demonstrated a low CCM burden throughout the brains with often large, focal areas of hyperintensity (Figure 5 D). We sectioned the brains in the regions of the focal hyperintensities, stained the endothelium with anti-CD31 antibodies, and visualized the autofluorescence of red blood cells. The presence of red blood cells in the brain parenchyma surrounding CCMs was indicative of acute hemorrhage (Figure 5 H). The amount of extralesional blood was quantified and reported as a percentage of the total lesion volume measured by microCT (Figure 5 B). Sixty percent of LPS injected *Pdcd10*^*BECKO/KO*^ mice (3/5) developed acute hemorrhage (Figure 5 B). One *Pdcd10*^*BECKO/KO*^ mouse injected with LPS was found dead 24 hours after LPS injection. This mouse is included in the analysis (open circle Figure 5B) and the entire characterization is shown in Supplemental Figure 3. We injected two cre-recombinase negative, littermate control genotypes (*Pdcd10*^*Flox/KO*^ and *Pdcd10*^*Flox/WT*^) with LPS and did not observe any CCM formation or bleeding from the cerebral vessels (n=4) (Figure 5 B, N). Thus, the acute hemorrhage induced by LPS in this *Pdcd10*^*BECKO/KO*^ model is specific to CCMs rather than a general response of the cerebral vasculature to LPS. In the present study, 60% (3/5) of the mice injected with LPS exhibited acute hemorrhage. This is the first model of acute CCM hemorrhage. Acute CCM hemorrhage is a life-threatening, medical emergency and this model provides the first system to test the ability of therapies to prevent or stabilize acutely bleeding CCMs.

## Discussion

Current CCM treatment consists of surgical removal or symptom management. There is no approved pharmaceutical therapy to treat the etiology or associated bleeding of this disease. While surgical intervention can be curative for patients with the sporadic form of the disease and a solitary CCM, these neurosurgical procedures are invasive and contain significant risk of associated morbidity and mortality. Not all CCM patients are candidates for surgery due to anatomic location of the lesion, with brainstem and deep lesions being particularly problematic. Thus, there is a significant need for a robust medical therapy to treat CCM lesion burden, but also hemorrhagic sequelae. Preclinical therapeutic studies require both strong drug candidates and animal models that more faithfully recapitulate important features of the human disease. Herein we expanded the repertoire of CCM mouse models by creating novel, inducible CCM3 mouse models of different stages of the human disease: 1) delayed formation of single cavern CCMs, 2) chronic CCM hemorrhage, and 3) acute CCM hemorrhage.

The inducible CCM mouse models have been an invaluable tool for mechanistic discoveries of CCM pathobiology but have a limited ability to test therapeutics in prolonged drug studies. The limitation of the inducible model arises from the severe CCM burden that leads to lethality near weaning, and lesions often lack associated inflammatory cell infiltrates and bleeding, which are hallmark features of the human disease. The rare exceptions to the early lethality include two studies with CCM1 [40] and CCM2 [32] inducible models that were able to complete drug studies lasting several months. The short treatment window for the vast majority of the inducible mouse models presents a challenge for administering therapeutics. A potentially confounding variable in the inducible models is the association of CCM formation with the developmental angiogenesis that continues after birth in the cerebellum. The CCMs that develop in these inducible models occur nearly exclusively in the cerebellum, suggesting a strong sensitizing role of angiogenesis. Inhibition of vascular endothelial growth factor (VEGF) signaling with SU5416, a VEGFR2 specific antibody, in an inducible CCM1 mouse model reduced CCM formation and hemorrhage [40]. By contrast, an exploratory biomarker study found plasma levels of VEGF to be lower in CCM patients who had a hemorrhage in the past 3 months when compared to CCM patients without hemorrhage [68]. Thus, the role of angiogenesis in CCM formation and hemorrhage in the human disease, as well as how it contributes to the phenotype of inducible mouse models remains unclear. We temporally separated developmental angiogenesis and CCM formation in the inducible *Pdcd10*^*ECKO*^ mouse model. We delayed *Pdcd10* deletion to P6 and observed CCM formation beginning at approximately P28, well after developmental angiogenesis has concluded. While the first CCMs in the *Pdcd10*^*ECKO*^ model develop in the cerebellum, the later CCMs develop throughout the entire brain in a pattern much more like the human disease. A more representative model of CCM disease will enable therapeutic studies that yield results with a greater likelihood to translate to human studies.

A robust CCM burden develops in the *Pdcd10*^*ECKO*^ model by P56 to enable relatively short-term drug studies, as opposed to several-month-long studies, to determine the ability of proposed therapeutics to impact CCM formation. When compared to the genetically sensitized mouse models [55], this inducible model is significantly more efficient at generating mice to enroll in studies; both the average litter size (7 versus 3) and the percentage of pups with the desired genotype (50% versus 17%) is greater in the inducible model. This inducible model is also able to develop a robust CCM burden in a fraction of the time that is needed for the genetically sensitized model. The tradeoff for developing a robust CCM burden in weeks, rather than months, is that the CCMs in the inducible model to not develop into mature, multicavernous and hemorrhagic lesions. We utilized the strengths of the *Pdcd10*^*ECKO*^ model to test the ability of fasudil, tempol, vitamin D_3_, and a triple therapy combination of these drugs to reduce single cavern CCM formation. As monotherapies, fasudil and tempol both trended towards a reduction of CCM burden and tempol demonstrated a statistically significant reduction in CCM burden. We then combined fasudil, tempol, and vitamin D_3_ as a combination therapy to see if additive or synergistic effects could be elicited. The triple therapy did not reduce the formation of CCMs. We attribute the modest results in our *Pdcd10*^*ECKO*^ model to 1) the known moderate abilities of each therapy to reduce CCM formation, 2) a difference in the CCM genes deleted in the mouse models, and 3) phenotypic variation of total lesion burden in the mouse model. Furthermore, we were not able to assess treatment effects previously shown on lesion maturation and hemorrhage in other models [30, 54, 55] in the *Pdcd10*^*ECKO*^ model. Our study highlights the limited ability of the studied drugs to reduce the CCM burden in a CCM3 model and the need for a more robust therapy.

There is a growing body of clinical evidence that CCM disease due to loss of *PDCD10* is much more aggressive than that from either *KRIT1* or *CCM2* loss [69-71]. KRIT1, CCM2, and PDCD10 also have distinct cellular roles and signaling pathways [52]. Given these differences at the clinical and molecular level, it would not be surprising to see a difference in CCM response to therapy depending upon the underlying genetic mutation of the malformation. The lack of a treatment effect in our *Pdcd10*^*ECKO*^ model with therapies that have shown an effect in CCM1 and CCM2 models supports the hypothesis of differential responses to therapy based upon which CCM gene is mutated.

Measuring the ability of drugs, with known modest effects, to treat CCMs is further compounded by the phenotypic variability inherent in any animal model of disease. The age at which *Pdcd10* is deleted plays a significant role in the severity and onset of CCM development. The CCM phenotype in the inducible model appears to be exquisitely sensitive to the amount of developmental angiogenesis occurring in the neonatal brain. Thus, variation in CCM burden in the *Pdcd10*^*ECKO*^ model may have occurred due to the variation of the exact age of the mice when *Pdcd10* deletion was induced. Because the precise time when an animal is born is almost always unknown, P6 animals might vary in age by as much as 12 hours. Therefore, the variability across different litters in the precise age of pups when *Pdcd10* was deleted likely translated into variability of the CCM burden of mice from different litters. We attempted to minimize the effect of the litter-to-litter variability by randomly enrolling pups from every litter into different treatment groups. The CCM research community continues to search for a robust therapy that has a strong effect that can easily be detected despite the phenotypic variation present in all drug studies with animal models.

Chronic CCM hemorrhage is a hallmark feature of the human disease for which few animal models exist for therapeutic testing [53, 51, 52, 58]. Very few studies have been able to use an inducible CCM mouse model to study a chronic CCM hemorrhage, none of which have been with the more aggressive CCM3 mouse model [40]. One reason for the paucity of studies measuring chronic hemorrhage is the need to conduct experiments over several months for the chronic hemorrhage phenotype to develop. We developed an inducible CCM3 mouse model of chronic CCM hemorrhage by restricting *Pdcd10* deletion to the cerebral vasculature, as opposed to *Pdcd10* deletion throughout the systemic vasculature with a pan-endothelial cre recombinase. Restricting *Pdcd10* deletion to the brain vasculature eliminated the systemic pathology observed with global endothelial cell deletion of *Pdcd10* (Supplemental Figure 1). Elimination of the systemic diseases, particularly within the gastrointestinal tract, significantly improved the viability of the mice and enabled the induced CCMs to develop into multicavernous lesions with chronic hemorrhage at postnatal day 121 (4 months). This inducible CCM3 model with chronic hemorrhage provides another model for the CCM research community to design therapeutic studies targeting this critical stage of the human disease and associated bleeding.

During preparation of this manuscript, another group developed a similar mouse model by restricting *Ccm2* deletion to the brain vasculature[72]. As observed in this study, their model also developed CCMs with chronic hemorrhage several months after *Ccm* gene deletion and suggests a reproducible phenotype can be induced in CCM1, CCM2, and CCM3 mouse models.

Acute CCM bleeding is a medical emergency. We describe here the first mouse model of acute CCM hemorrhage by injecting mice with existing CCMs with an environmental stimulus. We selected lipopolysaccharide (LPS) as the environmental sensitizer based upon the recent discovery of LPS, from Gram-negative bacteria in the gut, as a significant sensitizer of CCM disease [21]. We injected a single, sub-lethal dose of LPS and induced acute hemorrhage in 60% (3/5) of the mice. This model supports the previous findings of LPS as a significant sensitizer for CCMs and suggests that additional sources of bacterial LPS beyond the gut may contribute to CCM pathology. Similarly, additional TLR4 ligands may play a role in acute CCM exacerbation [73]. This model may be used to either administer a prophylactic therapy to prevent acute hemorrhage or administer a rescue therapy to stabilize a CCM after it begins to hemorrhage.

The new models we have generated provide a set of inducible CCM3 mouse models that will enable researchers to test drugs targeting specific stages of CCM pathology. It is unlikely that a single therapy will be a panacea for CCM disease: blocking CCM formation, preventing chronic hemorrhage, and preventing or stabilizing acute hemorrhage. It is much more likely that different therapies will be efficacious for different stages of disease and in malformations with different CCM gene mutations. These new models provide unprecedented control in designing preclinical studies to identify drugs to nominate for human clinical trials.

## Acknowledgements

The authors would like to thank Drs. Wang Min, Marcus Fruttiger, and Markus Schwaninger for generously providing transgenic mice used in this work. We would also like to thank Drs. Mark Kahn and Mark Ginsberg for helpful discussions.

## Funding sources

This work was supported by the National Institutes of Health (P01 NS092521 to D.A.

Marchuk and I.A. Awad, F30 HL140871 to M.R. Detter, and T32 GM007171), the Fondation Leducq (17 CVD 03 to D.A. Marchuk), and the American Heart Association (18PRE34060061 to M.R. Detter).

---

**Supplemental Figure 1.**
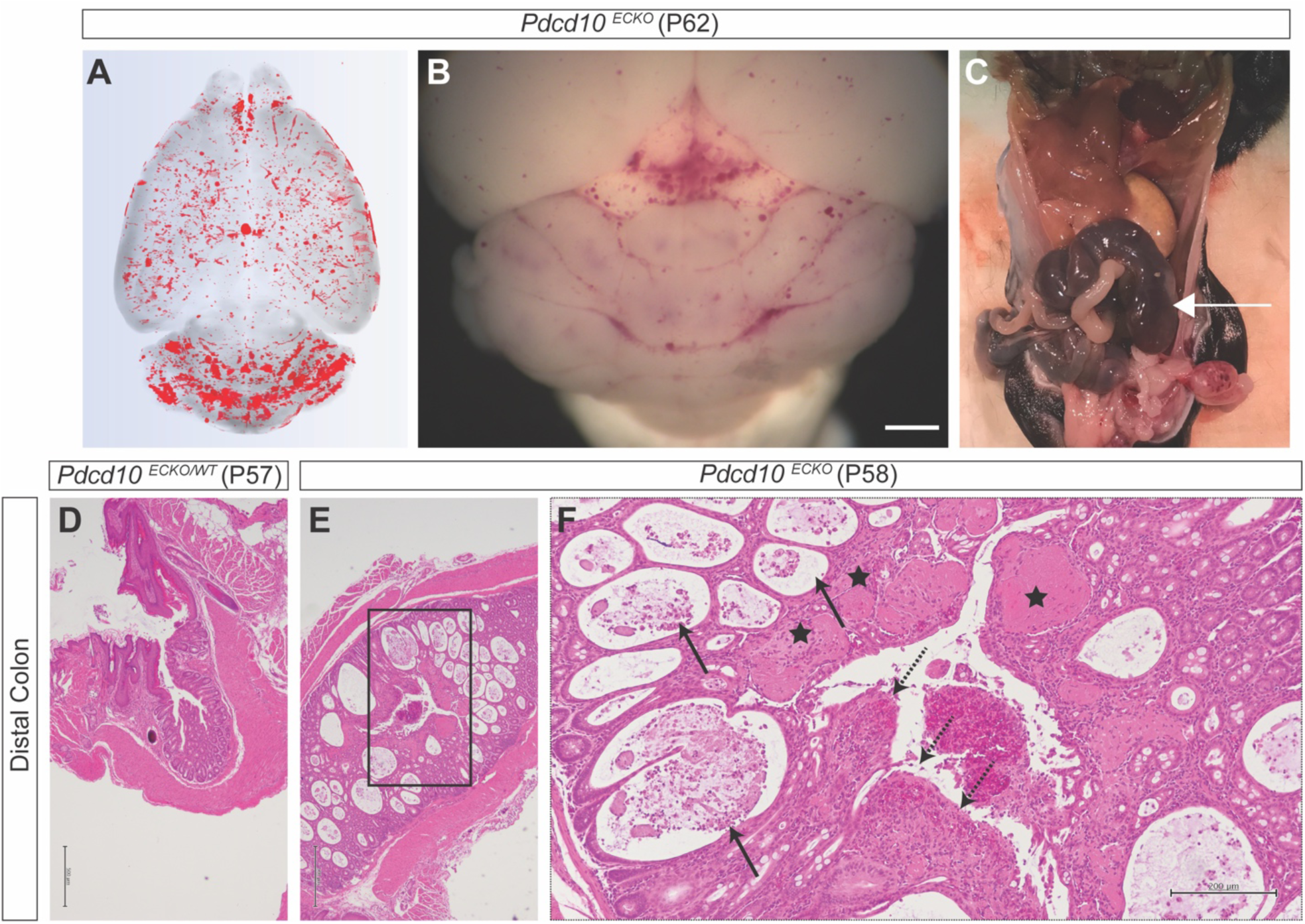
Deletion of *Pdcd10* throughout systemic endothelium leads to gastrointestinal pathology. A,B) MicroCT brain scan and gross brain image of a symptomatic *Pdcd10*^*ECKO*^ mouse euthanized on P62 (scale bar: 1mm). C) Gross image of the abdomen revealed ischemic bowels (arrow). D) Hematoxylin and eosin (H&E) stain the distal colon of a control mouse (*Pdcd10*^*ECKO/WT*^) at P57 demonstrating a normal colonic epithelium (scale bar: 500μm). E) H&E staining of the distal colon of a *Pdcd10*^*ECKO*^ mouse (Scale bar: E: 500μm) F) Higher magnification of E demonstrating dilated lamina propria micro-vessels (stars), crypt abscesses (solid arrow), and epithelial erosion with granulation tissue formation (dotted arrow) (Scale bar: 200μm).

**Supplemental Figure 2.**
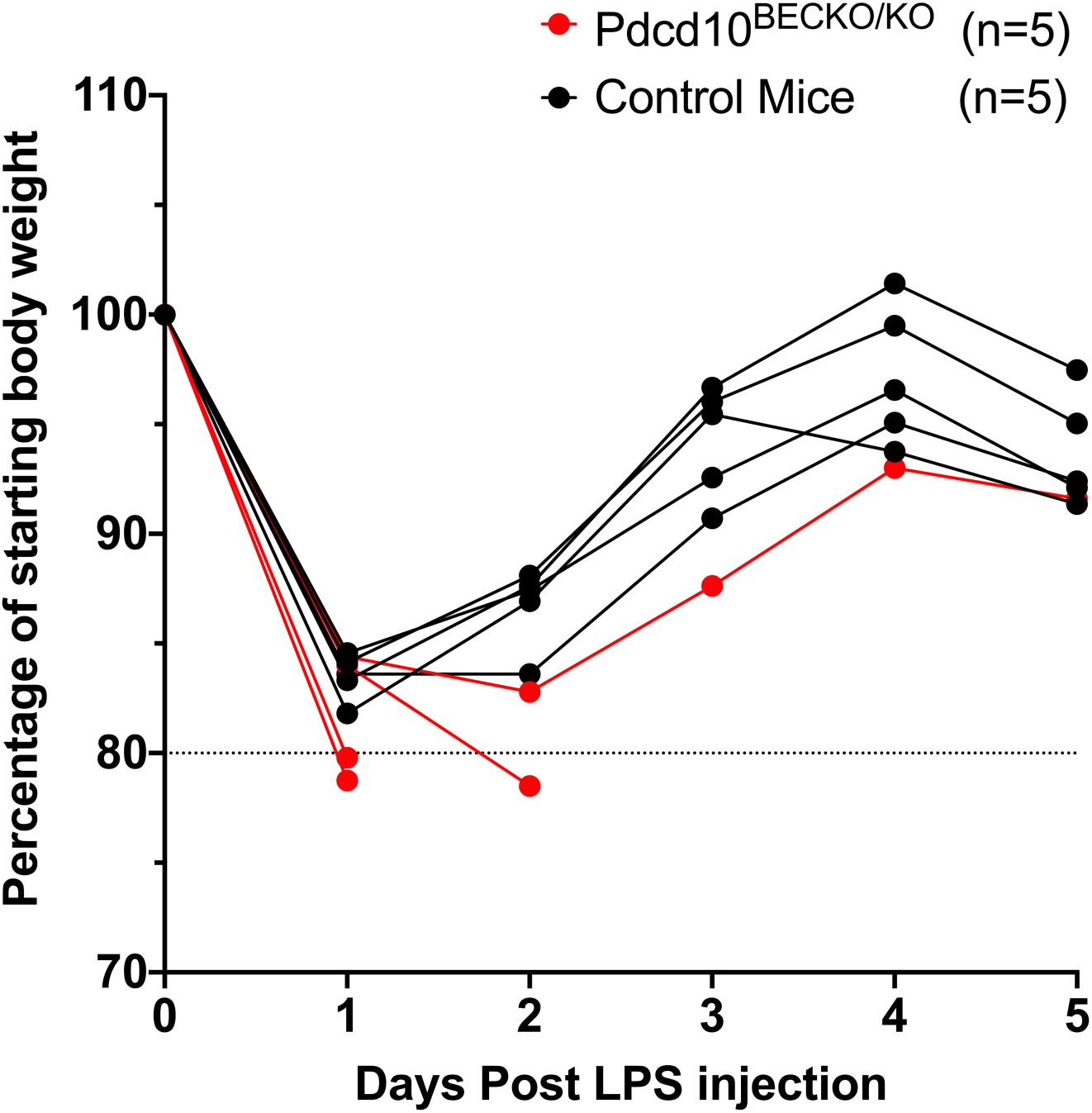
A single, sub-lethal dose of lipopolysaccharide (LPS) results in weight loss within 24 hours, reflecting an acute inflammatory response. The mice which did not lose >20% of their starting body weight, a humane endpoint for the study, recovered to their starting body weight by five days post LPS injection.

**Supplemental Figure 3.**
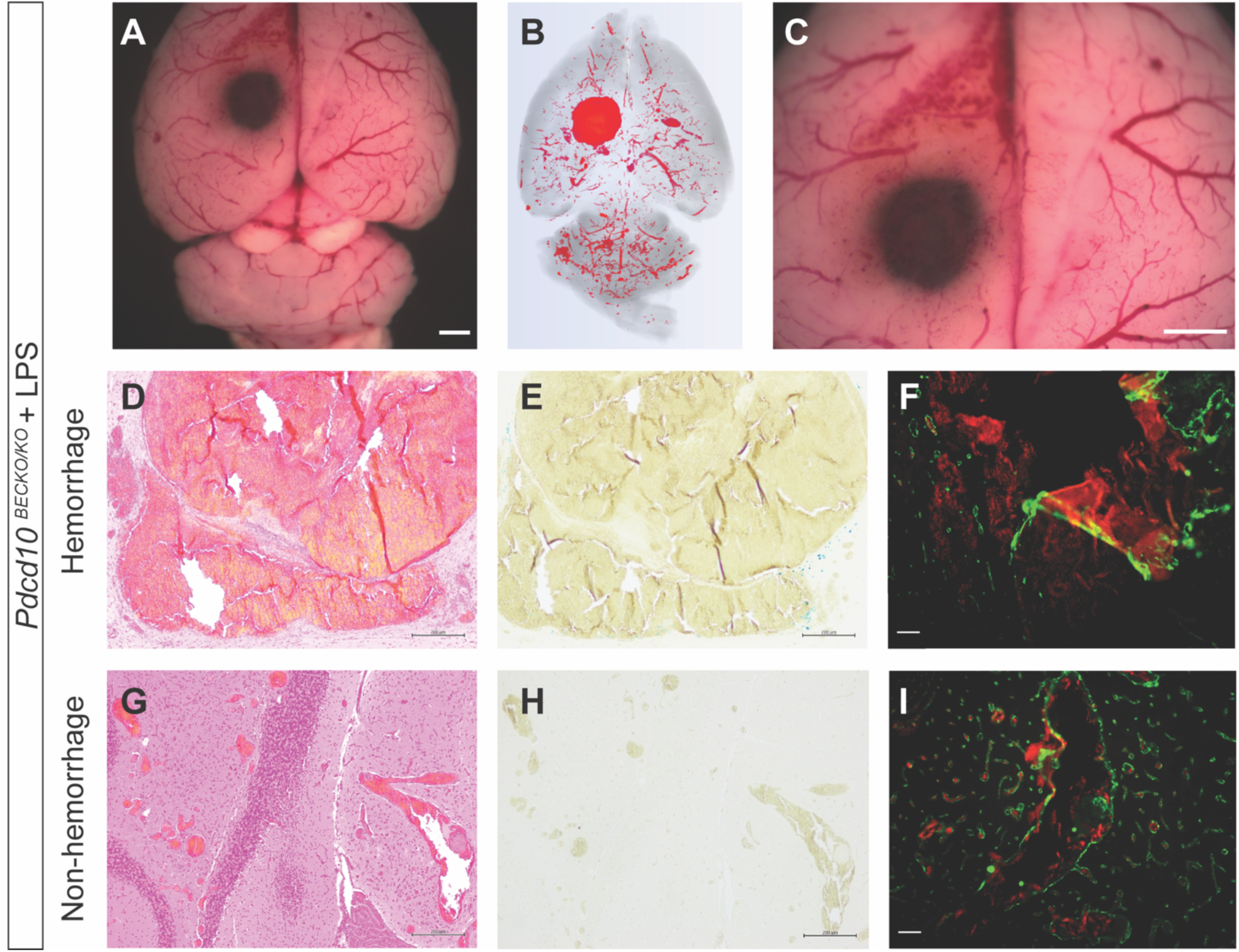
Histology of a *Pdcd10*^*BECKO/KO*^ mouse found dead 24 hrs after lipopolysaccharide (LPS) injection. A) Gross brain image with a large area of apparent hemorrhage in the left cerebral hemisphere (scale bar: 1mm). B) 3D rendering of the full brain microCT scan demonstrating the area of focal hyperintensity in the left cerebral hemisphere. C) Magnified view demonstrating a well circumscribed area of hemorrhage (scale bar: 1 mm). A hemorrhagic brain region of this mouse was analyzed with D) hematoxylin and eosin (H&E), E) Perls’ Prussian blue dye, and F) CD31-labeled endothelium (green) demonstrating autofluorescence (red) of red blood cells hemorrhaging from the CCMs into the brain parenchyma. A non-hemorrhagic brain region of this mouse was also analyzed with G) H&E, H) Perls’ Prussian blue dye, and I) CD31 labeling of endothelium with red blood cells remaining within the CCM lumen. (D,E,G,H scale bars: 200μm, F,I scale bars: 50μm).

